# Development and Application of Equilibrium Thermodynamic Models Describing Coupled Protein Oligomerization and Ligand Binding

**DOI:** 10.1101/068577

**Authors:** Martin L. Rennie, Richard L Kingston

## Abstract

The association and dissociation of protein oligomers is frequently coupled to the binding of ligands, facilitating the regulation of many biological processes. Equilibrium thermodynamic models are needed to describe the linkage between ligand binding and homo-oligomerization. These models must be parameterized in a way that makes physical interpretation straightforward, and allows elaborations or simplifications to be readily incorporated. We propose a systematic framework for the equilibrium analysis of ligand-linked oligomerization, treating in detail the case of a homo-oligomer with cyclic point group symmetry, where each subunit binds a ligand at a single site. Exploiting the symmetry of the oligomer, in combination with a nearest-neighbors approximation, we derive a class of site-specific ligand binding models involving only four parameters, irrespective of the size of the oligomer. The model parameters allow direct quantitative assessment of ligand binding cooperativity, and the influence of ligand binding on protein oligomerization, which are the key questions of biological interest. These models, which incorporate multiple types of linkage, are practically applicable, and we show how Markov Chain Monte Carlo (MCMC) methods can be used to characterize the agreement of the model with experimental data. Simplifications to the model emerge naturally, as its parameters take on extremal values. The nearest-neighbors approximation underpinning the model is transparent, and the model could be augmented in obvious fashion if the approximation were inadequate. The approach is generalizable, and could be used to treat more complex situations, involving more than a single kind of ligand, or a different protein symmetry.

**Author Summary:** The assembly and disassembly of protein complexes in response to the binding of ligands is a ubiquitous biological phenomenon. This is often linked, in turn, to the activation or deactivation of protein function. Methods are therefore needed to quantitate the linkage or coupling between protein assembly and effector binding, requiring the development of mathematical models describing the coupled binding processes. As proteins usually assemble in a symmetric fashion, the nature of any symmetry present has to be considered during model construction. We have developed a class of models than can effectively describe the coupling between effector binding and protein assembly into symmetric ring-like structures of any size. Despite the relatively complex mathematical form of the models, they are practically applicable. Markov Chain Monte Carlo methods, a form of Bayesian statistical analysis, can be used to analyze the fit of the models to experimental data, and recover the model parameters. This allows the direct assessment of the nature and magnitude of the coupling between effector binding and protein assembly, as well as other relevant characteristics of the system.

## Introduction

Proteins function by participating in chemical processes such as ligand binding, catalysis, conformational switching and oligomerization. Complexity arises when a single protein participates in multiple processes that depend upon one another. Interdependence of this kind has been termed linkage [1], coupling [2], or cooperativity [3,4] in the biological literature. Linkage of chemical processes is of cardinal importance, as it allows for the effective regulation of biological systems [5]

This paper concerns the linkage between protein homo-oligomerization and the binding of ions and small molecules, a ubiquitous biological phenomenon [6,7]. A prominent example is the zinc dependent-oligomerization of insulin. Insulin is stored as an inactive hexamer in pancreatic cells in the presence of elevated zinc concentrations. When released into the serum, insulin dissociates to form an active monomer, in response to the decrease in both zinc and proton concentrations [8]. Only monomeric insulin is capable of receptor binding and signaling pathway activation [9]. Quantitative analysis of such “assembly-binding” linkage is needed to advance basic biology. It is also needed when protein-protein interactions are targeted for therapeutic purpose [10], or when small molecules are used to manipulate the assembly of proteins into functional nano-materials [11-13].

Equilibrium studies are central to the quantitative analysis of coupled oligomerization and ligand binding. This approach requires an equilibrium thermodynamic model of the system, the collection of suitable experimental data, and a method of estimating the model parameters based on the data. These are the points we consider in this paper. Equilibrium thermodynamic models enumerate the discrete energetic states of the protein, and the equilibrium constants - or equivalently standard Gibbs energies - that determine the population of those states. When applied to ligand binding, models can be developed at different levels of detail (Fig. 1). Stoichiometric binding models consider states with the same total number of ligands bound to be equivalent. In contrast, site-specific binding models differentiate between the possible arrangements of bound ligand among the sites [14-16]. A stoichiometric model can always be derived from a site-specific model, however the converse is not true. In this sense site-specific models are more fundamental than stoichiometric models, and they are prerequisite for explaining binding phenomena at the molecular level.

**Figure 1.**
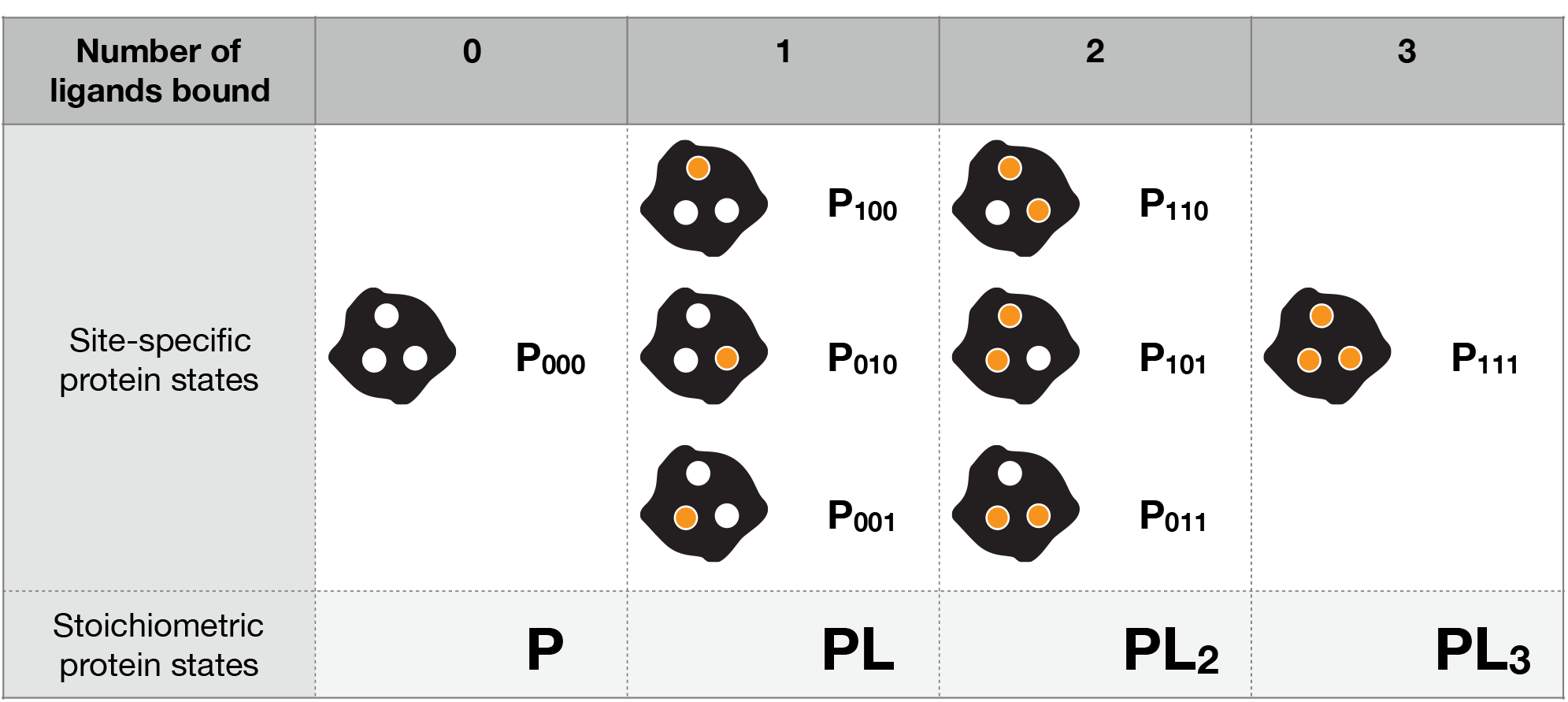
Stoichiometric versus site-specific ligand binding models. The equilibrium thermodynamic states of a protein that binds multiple ligands can be specified in terms of the total number of ligands bound (a stoichiometric model), or in terms of the possible arrangements of bound ligand (a site-specific model). The models developed in this paper are site-specific.

Unfortunately, the parameters of a site-specific model are often difficult to determine from experimental data. This is because many physical techniques for studying ligand binding do not differentiate between sites, and report only on the number of ligands bound. Unique recovery of the parameters of a general site-specific model is impossible from such data. Consequently, much of the quantitative analysis of protein ligand interactions has been performed using stoichiometric models. However site-specific models can be retained if they are suitably simplified, which can be achieved by incorporating additional information, or making reasonable approximations. Symmetry is one of the key features of proteins that can be exploited here. Homo-oligomeric proteins almost always have point group symmetry [17-20], the simplest form of which is cyclic symmetry. Symmetry implies energetic equivalence between many of the possible arrangements of bound ligand among the sites on the oligomer. In this paper, we demonstrate how incorporation of symmetry, combined with suitable approximations, results in site-specific models that describe ligand binding to dissociable protein oligomers, and can be usefully fit to experimental data.

The detailed formulation of site-specific ligand binding models requires that the parameterization of the model be carefully considered. While the number of parameters required for an exact equilibrium thermodynamic model is unambiguous, there is no unique way to assign equilibrium constants, nor any accepted convention for doing so. The use of alternative parameterizations complicates comparisons between different studies, and some methods of parameterizing complex binding models lead to mathematical intractability, or difficulties in interpretation. At least two systematic methods to parameterize equilibrium binding models have been proposed which explicitly account for the possible dependence between binding events [4,16,21,22], though these have not been widely employed in practice. Both schemes are hierarchical in nature, and describe ligand binding in terms of basal equilibrium constants for each site, modulated by interaction parameters of increasing order, which account for ligand binding cooperativity. If the sites operate independently, the interaction parameters vanish, and the models readily simplify. The difference between approaches arises in the definition of the interaction between three or more sites. Adoption of a systematic hierarchical parameterization scheme is critical in formulating a model that can be readily interpreted, and suitably simplified in either the presence or absence of linkage effects.

Irrespective of the exact form of the model, parameter estimation for assembly-binding equilibrium models requires measurement of ligand binding and/or protein oligomerization as a function of total ligand and protein concentration. Often ligand binding data is collected under conditions where one oligomeric state dominates. Alternatively oligomerization data is collected in the absence of ligand and in the presence of saturating amounts of ligand. This allows some of the component equilibria to be characterized in isolation, using simple models that neglect linkage. Non-linear least squares techniques are typically used to perform model fitting. However, when the objective is to quantitate linkage, a more direct approach is to freely vary the concentration of both protein and ligand, and fit the data globally to a model that incorporates the linkage effect. Such models will have a relatively complex mathematical form, requiring more powerful parameter estimation techniques. Here we explore the use of Markov Chain Monte Carlo (MCMC) methods for model fitting, which are a form of Bayesian inference [23-25]. MCMC methods provide the posterior probability distributions of the model parameters given the data, which is particularly useful for fitting complex models to limited experimental data. Even if reliable point estimates for the model parameters cannot be obtained, it is often still possible to learn something about their bounds.

The simplest example of the phenomenon we are investigating - a monomer-dimer system with a single ligand binding site per subunit - has been well studied at the theoretical level [26-32]. Gutheil has discussed the systematic parameterization of an equilibrium thermodynamic model for this case [32]. Although there are many experimental studies of such systems (for recent examples see [33-35]), we are aware of none that have employed a systematic and non-redundant parameterization of the underpinning equilibrium thermodynamic model. Furthermore, in this simplest of cases, stoichiometric and site-specific ligand binding models are essentially equivalent, hiding the complexities that emerge when large oligomers are studied. Models for oligomerizing systems of arbitrary complexity have been proposed [6,36,37], however they are stoichiometric with respect to ligand binding, and not practically applicable due to the large number of model parameters.

In this paper we detail a systematic quantitative method for the equilibrium analysis of coupled ligand binding and protein oligomerization. The specific problem treated is that of dissociable protein oligomers with cyclic point group symmetry, where each subunit can bind ligand at a single site. However the approach could be readily extended to encompass more complex situations. We develop the model in a stepwise fashion. Beginning with a general site-specific ligand binding model, appropriately parameterized, we apply cyclic symmetry restrictions. We then make an Ising approximation, assuming each subunit only senses the state of its immediate neighbors. Finally we extend the model to allow for dissociation of the oligomer into its constituent subunits. This results in a set of binding models governed by only four parameters, irrespective of the size of the oligomer. The model is exact for the monomer-dimer case. The parameters of the model have a clear physical meaning, and quantitate ligand binding cooperativity and the linkage between ligand binding and oligomerization. When developed mathematically, we show that the models describe literature data well, and that MCMC techniques can be used to estimate the parameters of such site-specific models from limited binding data. Finally, we use the models for theoretical exploration of the ligand binding problem, and demonstrate how biologically interesting scenarios can emerge in some extremal cases.

## Results

### Derivation of Equilibrium Thermodynamic Models

The models developed here assume that experiments are carried out at constant temperature and pressure on ideal associated solutions [38], in which the only departures from ideality arise from the chemically specific and reversible interactions between protein and protein, and protein and ligand. For simplicity only binding of a single type of ligand to the protein is considered, though the approach is readily generalizable.

#### General site-specific ligand binding model

To begin, consider a general site-specific model for a protein with *n* ligand binding sites, where the sites are not identical, and may not be independent. The model enumerates all of the possible configurations of bound ligand, which are taken to be the thermodynamic states in which the protein can exist. The simplest cases, with two, three and four sites, are readily diagrammed (Fig. 2, Fig. 3A, Fig. S1A, respectively). Site-specific ligand binding models of this kind are of mostly theoretical interest, owing to the large number of parameters (2*^n^* - 1) required for their complete specification.

**Figure 2.**
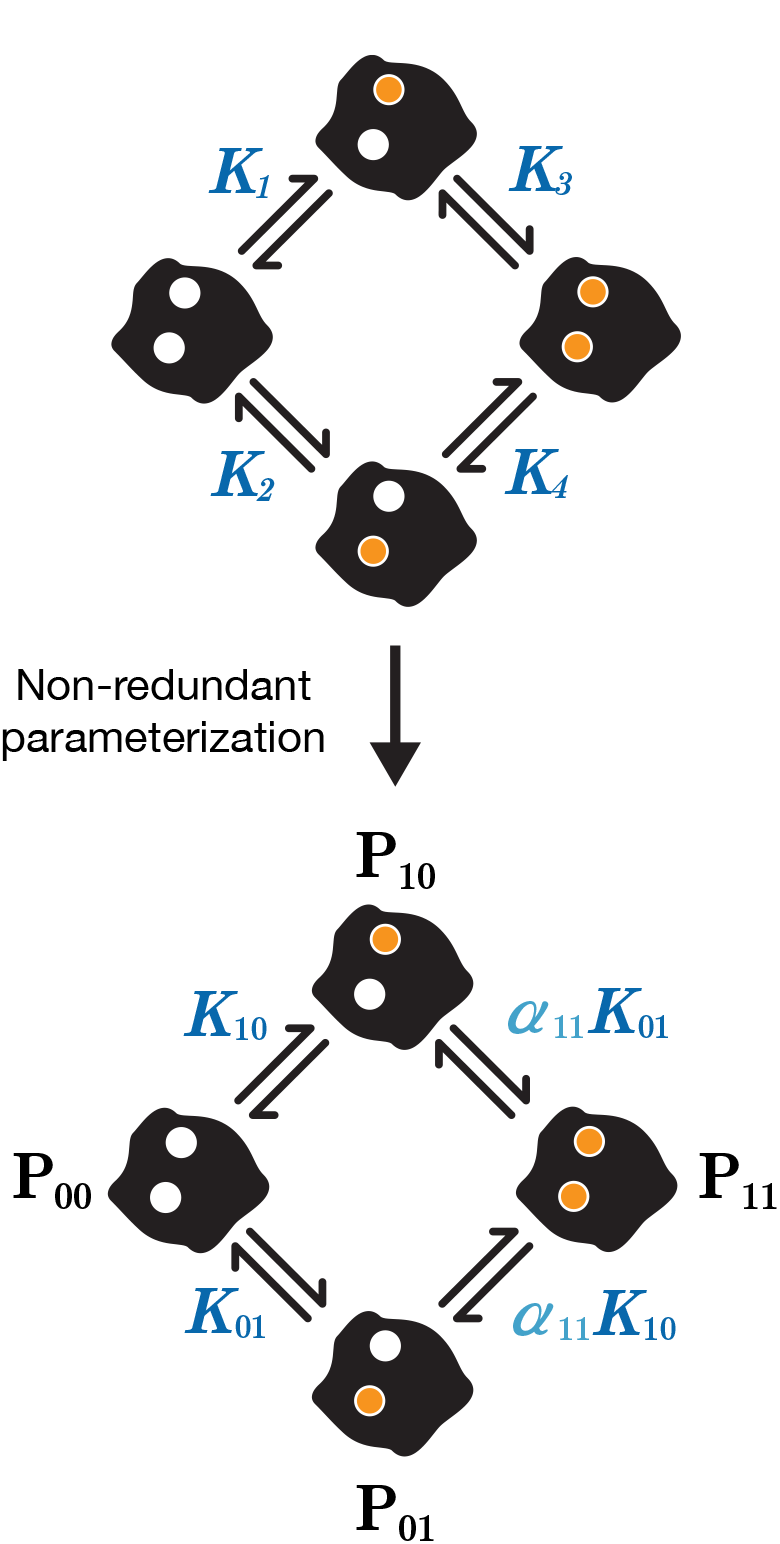
Parameterization of the general two-site ligand binding model. A simple approach to parameterizing equilibrium ligand binding models is to assign an equilibrium constant to each of the fundamental binding steps (top panel). However the presence of a thermodynamic cycle requires that *K*_1_*K*_3_ = *K*_2_*K*_4_, making the assignment redundant. The redundancy can be eliminated by definition of basal equilibrium association constants for binding to each site, as well as a pairwise linkage constant between the sites (bottom panel). A binary vector notation provides a systematic way to specify the states in which the protein can exist, and the parameters that govern the equilibria between states. See Fig. S2 for the equivalent parameterization in terms of Gibbs energies of reaction.

**Figure 3.**
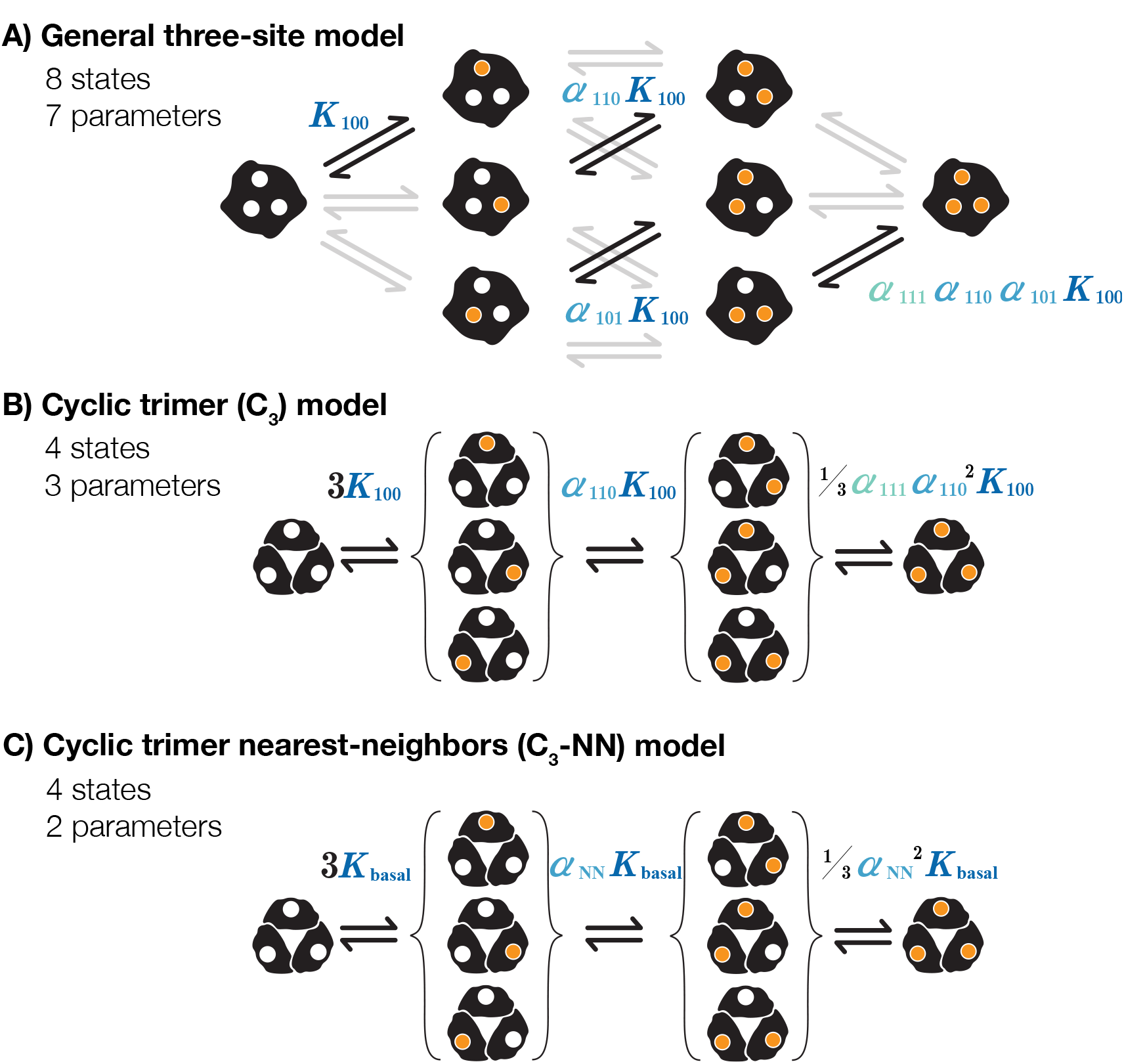
Schematic derivation of the C_3_-NN model. (A) The general site-specific ligand binding model is shown for a protein with three sites. Equilibria involving binding of ligand to site 1 are highlighted, to illustrate how they are governed by compound terms involving a basal binding constant and one or more linkage constants. A color gradient illustrates the hierarchal nature of the linkage constants. (B) Cyclic symmetry greatly simplifies the model reducing both the number of states and parameters. Symmetry equivalent states are bracketed. Imposition of cyclic symmetry results in the following formal parameter constraints: *K*_100_=*K*_010_=*K*_001_; *α*_110_=*α*_101_=*α*_011_. (C) The assumption that only adjacent subunits are coupled further simplifies the model. This nearest-neighbors approximation eliminates a single parameter (*α*_111_=1) from the cyclic model, however with larger numbers of sites, more parameters are eliminated (see Fig S1C for four-site case). For simplicity the parameters are relabeled (*K*_basal_=*K*_100_=*K*_010_=*K*_001_, *α*NN=*α*_110_=*α*_101_=*α*_011_).

The assignment of parameters governing the equilibria makes such models quantitative. The thermodynamic states can interconvert by binding or release of individual ligands and are therefore linked by a network of second-order equilibria (Fig. 2, Fig. 3A, Fig. S1A). An obvious way to proceed is to assign an equilibrium constant (or equivalently, a standard Gibbs energy of reaction) to each of these fundamental binding steps. However such a description is always redundant, involving more parameters then are needed to specify the equilibrium position of the system [15]. This parameter redundancy impedes both practical application and theoretical analysis.

This problem is addressed by a systematic and non-redundant parameterization previously proposed by Gutheil and McKenna [21]. In this parameterization, basal binding parameters quantify ligand binding to the individual sites of the unligated protein, and one or more additional parameters quantify the interaction between multiple sites (i.e. allow for linkage between sites). The “linkage parameters” constitute an additive hierarchy [21] (i.e. higher order linkage parameters represent the linkage additional to the the lower order linkages). Consider the two-site case (Fig. 2). Binding of ligand at the second site, given the first is occupied, can be described using a basal binding parameter modulated by the linkage parameter representing the pairwise (or second-order) interaction between the first and second sites. This modulation is multiplicative if the model is represented in terms of equilibrium constants (Fig. 2) and additive if represented in terms of Gibbs energies (Fig. S2) due to the logarithmic relationship between equilibrium constants and standard Gibbs energies of reaction. Adopting this parameterization, stepwise equilibria are governed by compound terms involving a basal binding parameter and one or more linkage parameters. When there is no interaction between sites, or when there is symmetry in the system, such that some of the sites are equivalent, the model is readily simplified. While the parameterization of Gutheil and McKenna is used here, related parameterizations have been described which differ slightly in the definition of the higher order linkage parameters [4,16,22].

In addition to a systematic parameterization, treatment of multi-site ligand binding requires a clear and consistent notation for the states the protein can exist in, and the parameters that govern the equilibrium position of the system. To represent the protein states a binary notation is adopted. An *n*-dimensional vector denotes the state of a protein possessing *n* ligand binding sites (cf [4,16,22]). Each vector element represents the ligation state of a site, 1 indicating the site is occupied, and 0 indicating the site is unoccupied. Hence P_000_ represents a three-site protein with no sites occupied, and P_101_ represents a three-site protein with the first and third sites occupied. When specifying the model in terms of equilibrium constants, K is used to denote basal association constants and α is used to denote linkage constants. When the corresponding terms are represented as standard Gibbs energies of reaction, they are denoted with Δ_r_G° and γ, respectively. Subscripts on these parameters indicate the sites they relate to. For example K_010_ denotes the basal association constant for binding to site two of three sites, while α_011_ denotes the pairwise linkage constant between sites two and three.

#### Ligand binding to oligomers with cyclic point group symmetry

Many protein oligomers have cyclic point group symmetry. We treat ligand binding to cyclic oligomers to illustrate how symmetry can be incorporated into the model. If each subunit of the oligomer can bind a single ligand, then the notation introduced above can be used without modification. Vectors now denote the ligation state of the *n* subunits of the oligomer, rather than the *n* sites on a single protein. The advantage of the vector notation is that it makes treatment of symmetry quite straightforward. The effects of symmetry on the model can be systematically analyzed using permutation matrices (see supporting information, Section S1).

By applying cyclic permutation, the symmetry equivalence of ligand configurations can be quickly assessed. Incorporation of symmetry results in a large reduction in the number of states, and therefore the number of parameters, when compared to the general ligand binding model (Table 1), and the introduction of degeneracies associated with sets of symmetry-equivalent states. For example, in the three-site case, imposition of cyclic symmetry equivalences the states *P*_110_, *P*_101_ and *P*_011_. They are no longer physically discriminable, and must be represented by a single macro-state. Without loss of generality, we can represent this triplet of symmetry equivalent states as 3*P*_110_. The imposition of symmetry also requires that the appropriate “statistical correction factors” are incorporated when specifying the stepwise equilibria, which account for the degeneracies introduced by symmetry (see [21,32]). The effects of incorporating cyclic symmetry are illustrated schematically for a three-site trimer and a four-site tetramer (Figs. 3B & S1B).

The problem of enumerating the non-equivalent states and their degeneracies, which is critical to the mathematical development of the models, is not entirely trivial. A point that only becomes apparent with four-fold and higher symmetry, is that states with the same number of ligands bound can be non-equivalent. For example, a cyclic tetramer with two sites bound (Fig. S1B), can exist in one of two different states, with differing degeneracies. The enumeration of all the distinct configurations can be achieved by inspection for low-order symmetry, or by the group theoretic technique of double coset decomposition for high-order symmetry [39]. For cyclic dimer to heptamer, the non-equivalent configurations of bound ligand and their associated degeneracies are detailed in Table S1.

**Table 1.**
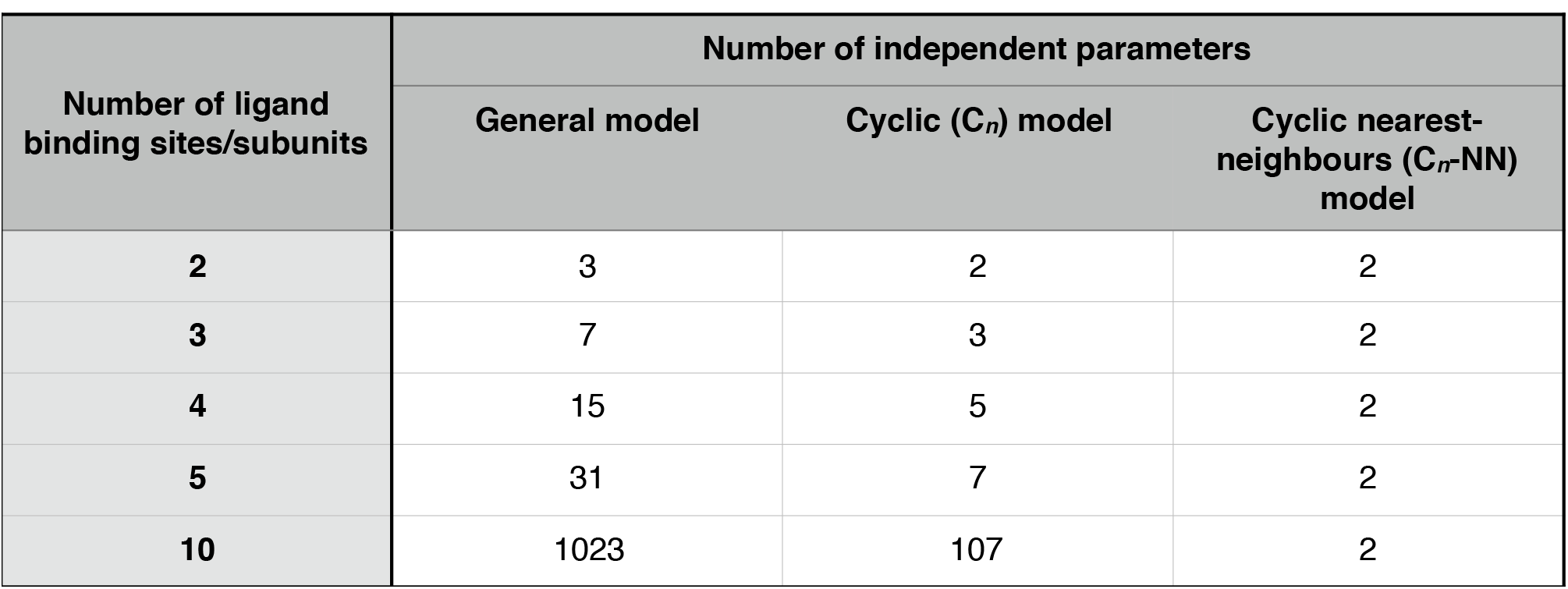
Number of parameters required for equilibrium thermodynamic models as a function of the number of ligand binding sites.

#### Nearest-neighbor (NN) approximation

Incorporation of cyclic symmetry dramatically reduces model complexity compared to the general case. However the number of model parameters still becomes very large as the size of the oligomer increases (Table 1), and some approximation is necessary for practical application. Models of multi-site ligand binding systems often incorporate only pairwise interactions between spatially adjacent subunits. This “nearest-neighbor” approximation has been applied in numerous contexts, one prominent example being lattice models of protein nucleic acid interactions [40]. Mechanistically, interaction between sites will often result from allosteric communication across the subunit interfaces of the oligomer. Most cyclic oligomers only have a single type of subunit-subunit interface due to the symmetry, and the presence of a central hollow pore. Hence it seems inherently reasonable to introduce a nearest-neighbor approximation for ring-like protein oligomers. For the cyclic dimer there is only a single nearest-neighbour and no approximation is involved.

Applying this approximation sets the pairwise linkage constants between non-adjacent sites to unity, together with all of the higher order linkage constants. That leaves a single nearest-neighbor (pairwise) linkage parameter and a single basal binding parameter in the model (Figs. 3C & S1C). The basal binding and nearest-neighbor linkage parameters are denoted K_basal_ and α_NN_ in terms of association equilibrium constants, or Δ_r_G°_basal_ and γ_NN_ in terms of Gibbs energies of reaction. The critical feature of these cyclic nearest-neighbor (C_n_-NN) models is that they have only two parameters regardless of the number of subunits in the oligomer (Table 1). These parameters will be estimable from even stoichiometric ligand binding data. Unlike the empirical Hill model [41] which also has two parameters, and is often employed to characterize binding cooperativity, C_n_-NN models have a solid theoretical grounding and the parameters have a straightforward molecular interpretation. Their application simply requires that the oligomeric state and symmetry of the protein is established, which is usually the case.

The C_n_-NN models are equivalent to those previously derived based on the Ising model [4,22,42-45]. However our derivation makes clear the relationship to the general ligand binding model, and the attendant assumptions and approximations involved in the derivation. Furthermore, the approach here is far more general, and readily adaptable to other situations involving different protein symmetries (cf [46]), the binding of additional ligands, or the presence of non-negligible higher-order interactions. One situation where the latter might occur is when we depart from the “small ligand” scenario. We briefly discuss this case in the supporting information, Section S2.

#### Oligomer Dissociation

Many protein oligomers undergo reversible dissociation into their constituent subunits at lower protein concentrations, which can be critical for their biological function. Oligomer association and dissociation can also be linked to ligand binding. We now consider how to elaborate the C_n_-NN models to incorporate these complexities.

We will assume that only the monomer and the cyclic oligomer are appreciably populated at equilibrium This model should be widely applicable, as equilibrium intermediates in the assembly of cyclic oligomers have seldom been detected experimentally [47-49]. Under this assumption only two additional protein states are introduced - the unligated and the ligated monomer. This is illustrated for the trimeric and tetrameric case in Figs. 4 and S3, respectively The presence of two additional states requires introduction of two additional model parameters. Again the parameterization requires careful consideration, and we adopt the approach of Gutheil [32]. In the dissociable case, the most convenient choice of “reference state” is the unligated monomer (in contrast to the non-dissociable case, where the most convenient choice is the unligated oligomer). Basal parameters quantitate the ligand binding and oligomerization of the unligated monomer. Ligand binding to the unligated oligomer is then described in terms of the basal binding to the monomer, modulated by the linkage between ligand binding and oligomerization (Figs. 4 and S3) [32]. This same linkage parameter is then systematically propagated through the other fundamental steps in the binding scheme.

**Figure 4.**
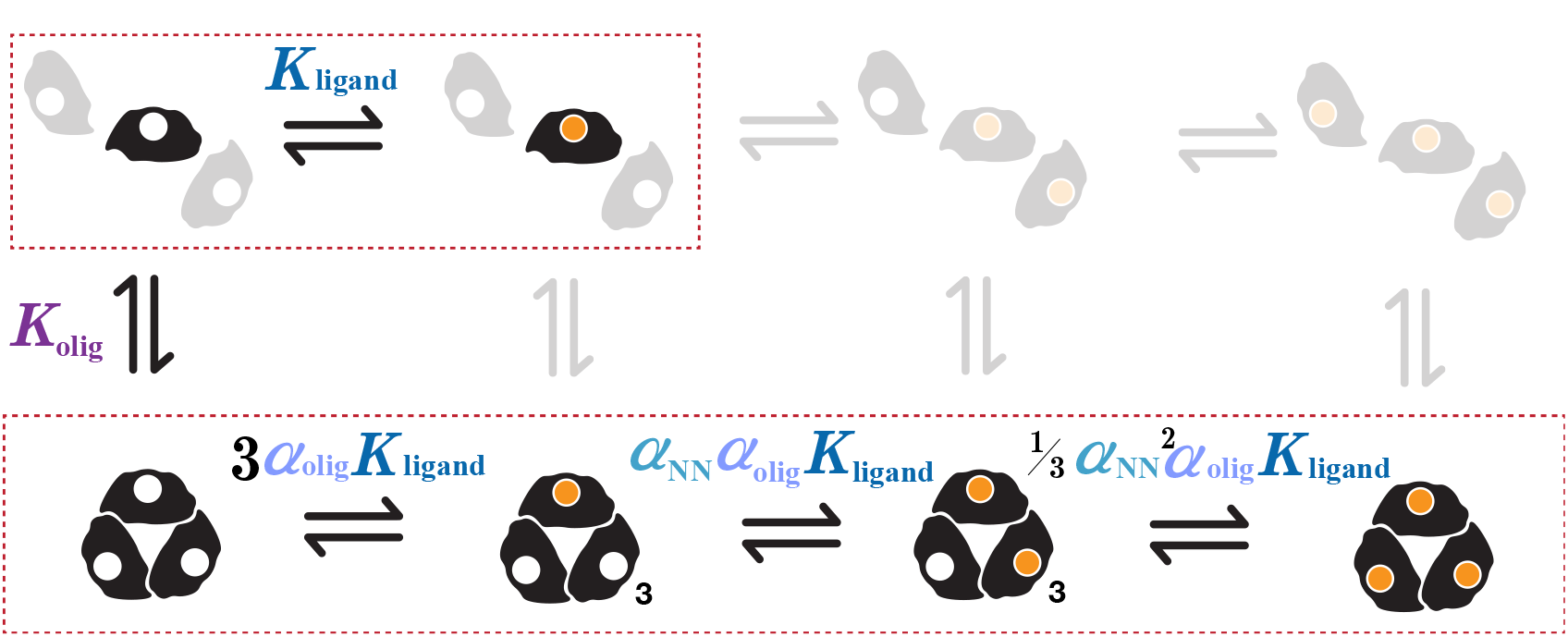
The dissociable C_3_-NN model. Extension of the C_3_-NN model (Fig. 3C) to include oligomer dissociation into constituent subunits introduces two monomeric states in addition to the trimeric states. For clarity only one representative of each symmetry equivalent state is shown, with a subscript denoting the associated degeneracy. Ligand binding equilibria are arranged horizontally and oligomerization equilibria are arranged vertically. Ligand binding equilibria of the monomer and trimer, respectively, are grouped by dashed boxes. From the highlighted equilibria, the terms governing all other steps can be deduced by consideration of the component thermodynamic cycles. See also Fig. S4, which shows the equivalent parameterization of the dissociable C_3_-NN model using standard Gibbs energies of reaction, and Fig. S3, which shows the dissociable C_4_-NN model.

In summary, when represented in terms of equilibrium constants, the four parameters governing the dissociable C_n_-NN models are: (1) a parameter specifying ligand binding to the monomer (K_ligand_) (2) a parameter specifying oligomerization of the unligated monomer (K_olig_), (3) a linkage parameter describing the coupling between oligomerization and ligand binding (α_olig_), and (4) a linkage parameter describing the coupling between ligand binding at adjacent sites (“nearest-neighbors”) on the oligomer (α_NN_). There is of course a completely parallel representation of the model in terms of standard Gibbs energies of reaction (see Fig. S4 for the trimeric case). This model is exact for the monomer-dimer system. For all higher symmetries it is valid if the nearest neighbors approximation holds, and there are no equilibrium intermediates between monomer and oligomer.

If the linkage parameters of the model can be experimentally determined, then the coupling present in the system is immediately accessible. Considering oligomerization and ligand binding, the sign and magnitude of γ_olig_ (=-RTlnα_olig_) quantitates the coupling between the two process. If γ_olig_ < 0 (or equivalently α_olig_ > 1) then they are positively coupled, if γ_olig_ > 0 (or equivalently α_olig_ < 1) then they are negatively coupled, and if γ_olig_ = 0 (or equivalently α_olig_ =1) then the two process are independent. In analogous fashion, the sign and magnitude of γNN (=-RTlnα_NN_) quantitates ligand binding cooperativity, just as it does for the non-dissociable case.

### Mathematical development of the dissociable C_n_-NN models

#### Specification of the binding polynomial

Development of the dissociable C_n_-NN model has proceeded schematically to make the basic form of the model clear, and to highlight the role of symmetry and the nearest neighbors approximation. However for practical or theoretical application, the model must be developed mathematically.

An equilibrium thermodynamic model of a binding process may be conveniently summarized via a “binding polynomial” (or partition function), from which useful expressions for many equilibrium properties of the system may be derived [1]. The key feature of the binding polynomial is that it represents the concentrations of all states, generally expressed relative to the concentration of some reference state. When modeling homo-oligomerization, it’s most convenient to neglect the normalization, and define the binding polynomial (Q) as a simple summation of the species concentrations [1]. For example, in the monomer-trimer case (dissociable C_3_-NN model, Figs. 4 and S4) it is given by:

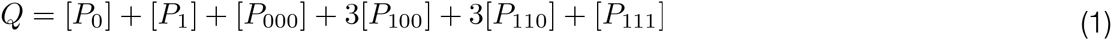

Using the definition of the equilibrium constant, we can re-write the concentration of each state in terms of the thermodynamic model parameters (K_ligand_, K_olig_, α_olig_, α_NN_), the free (unbound) ligand concentration ([*L*_free_]), and the concentration of a protein reference state, which we have taken to be the unligated monomer ([*P*_0_]).

Again, using the dissociable C_3_-NN case to illustrate, we have by inspection of Fig. 4

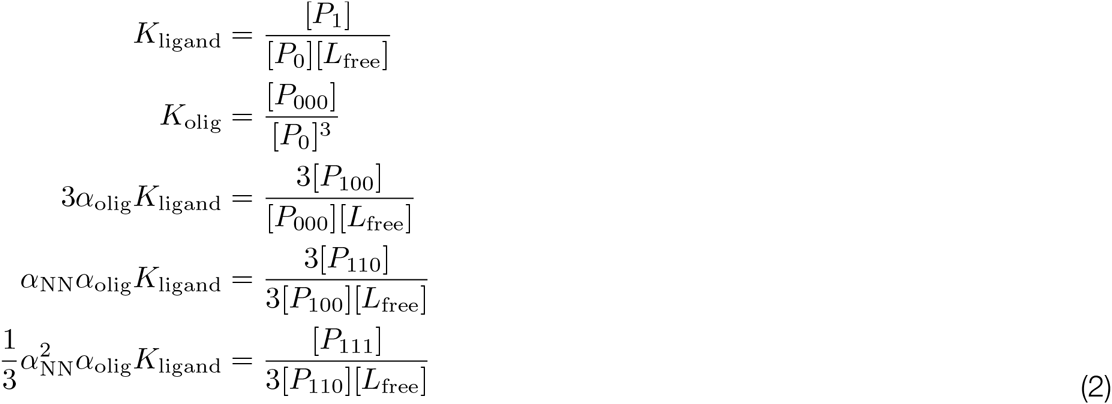

Using these equations we can rewrite the dissociable C_3_-NN binding polynomial (Eq. 1) as

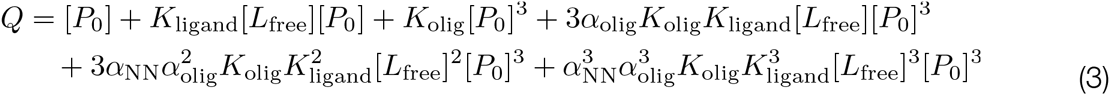

This approach can be straightforwardly extended to dissociable C_n_-NN models of any order, though it becomes quite laborious for large n. However, an alternative technique exists - the transfer-matrix method - that can be utilized to quickly determine the binding polynomial of even very large cyclic oligomers (supporting information, Section S3). The binding polynomials for some dissociable C_n_-NN models (n = 2,3,4,5) are given in Table S2. It is critical to appreciate that while the dissociable C_n_-NN models have only four parameters irrespective of oligomer size, the binding polynomials differ markedly with n.

With the binding polynomial specified, almost all of the mathematical tasks needed for model fitting and analysis can be readily handled. Most fundamentally, we need to be able to predict the concentrations of all states, given the thermodynamic model parameters, as well as the total ligand concentration ([L_Total_]) and total protein subunit concentration ([P_Total_]), which are generally the compositional variables under experimental control. The route to achieve this involves the mass balance equations for the protein subunits (Eq. 4) and for the ligand molecules (Eq. 5), both of which can be written concisely in terms of the (unnormalized) binding polynomial [1].

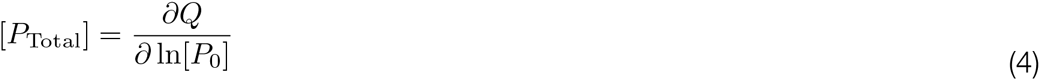

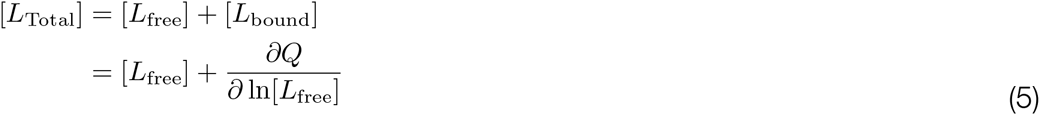

When the appropriate binding polynomial is inserted the mass balance equations readily simplify, using differential calculus, into polynomial equations. For example, for the dissociable C_3_-NN model these expressions have the explicit form:

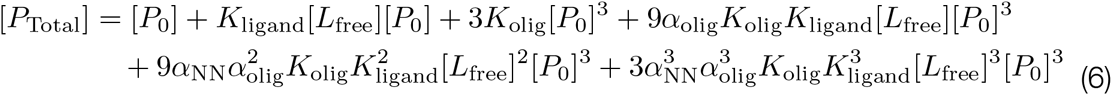

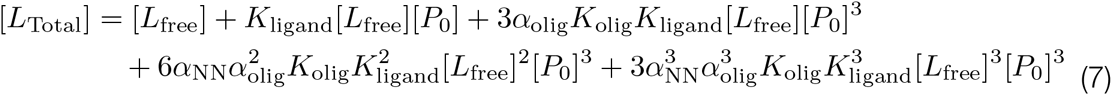

These constitute a pair of simultaneous equations written in terms of two unknowns (the free ligand [*L*_free_] and unligated monomer [*P*_0_] concentrations). While a solution for these quantities must exist, it cannot be expressed in closed form in even the simplest case (the dissociable C_2_-NN model). Hence the mass balance equations must be solved numerically, as we discuss below. With [*L*_free_] and [*P*_0_] determined (given estimates for the model parameters and the total protein and ligand concentrations) the concentrations of the remaining states are specified by the individual terms in the relevant binding polynomial. We note that when the ligand can be effectively buffered, and [*L*_free_] is under experimental control, only the mass balance equation for the protein subunits need be considered. In this special case, analytic solutions for [*P*_0_] can be obtained for n < 5.

The binding polynomial can be used to recover other properties useful for model fitting or analysis. For example, with an oligomerizing system, knowledge of the number-average, mass-average or Z-average molar mass of the protein is often required to interpret experimental measurements. These mean quantities can be straightforwardly computed from the molar mass of each state, and the corresponding terms in the binding polynomial.

### Practical application of the dissociable C_n_-NN models: General Aspects

The experimental study of coupled oligomerization and ligand binding will generally involve measuring one or more equilibrium properties as the total protein and ligand concentrations are varied. For example, the concentration of free ligand might be directly determined, or some physical observable measured that is responsive to ligand binding and/or protein oligomerization. Common examples of the latter are intrinsic protein or ligand fluorescence, heat change, or protein hydrodynamic properties. To assess the applicability of the dissociable C_n_-NN model and reliably determine all of its 4 parameters, measurements of several different types will be needed, which collectively monitor both ligand binding and protein oligomerization.

There are two aspects to mathematically modeling the resulting experimental data. The first problem is to describe the equilibrium thermodynamic behavior of the system. This is resolved using the dissociable C_n_-NN model. This model can describe the population of the protein states as a function of its 4 parameters (which are unknown, and of fundamental interest) and the total ligand and protein concentrations (which are known, and under experimental control). The second problem is to describe the relationship between the population of protein states and the experimental observables. Modeling of the experimental observables will involve additional physical parameters that are specific to the measurements being made. These parameters are also usually unknown, but of subsidiary interest. For clarity we term the parameters of the dissociable C_n_-NN model the thermodynamic parameters, and the parameters required to model the experimental observables the signal parameters. While the modeling of the experimental signal is important, and critical to successful practical application of any thermodynamic model, it is also entirely case-dependent, and not our primary concern in this paper.

With a complete model formulated, the problem of fitting it to the experimental data needs consideration. While non-linear least squares procedures are used extensively in simple ligand binding studies [50], and could be employed here, the reliability of the resulting point estimates for the model parameters is very difficult to assess. At present, Markov Chain Monte Carlo (MCMC) analysis [23-25] appears to be the best general method to overcome this problem, and analyze the fit of these relatively complex models. This form of Bayesian statistical analysis generates the posterior probability distributions for the model parameters, given the data, allowing straightforward identification of well-determined and ill-determined parameters. Posterior distributions for simple functions of the model parameters can also be determined using this method. For example, the equilibrium association constant for ligand binding to the “empty” oligomer might be of interest, which is proportional to the product of two parameters (α_olig_K_ligand_) of the dissociable C_n_-NN model (Figs. 4 and S3). Useful tutorials on the application of MCMC methods with biophysical examples can be found in [23,24] and many handbooks have been written (e.g. [25]).

We implemented an iterative MCMC algorithm for fitting dissociable C_n_-NN models to experimental data in Mathematica (Wolfram Research). At each iterate the mass balance equations (4 and 5) are solved numerically to yield the concentrations of all species, given the relevant total concentration of protein and ligand, and the current thermodynamic model parameters. Subsequently, the complete model for the experimental observables is evaluated, using the current signal model parameters. All model parameters were then updated using the Metropolis-Hastings algorithm [51,52], and the procedure iterated until the parameter distributions converged, as fully described in the supporting information, Section S5.

### Practical application of the dissociable C_n_-NN models: Specific Examples

To establish the practical utility of the dissociable C_n_-NN models, we re-analyzed experimental data from two previously published studies reporting ligand binding to dissociable protein oligomers. In re-analyzing the data we have used the dissociable C_n_-NN model to describe the equilibrium thermodynamic behavior of each system. The authors original model for the relationship between the experimental observables and the system composition was retained, facilitating comparison of the results.

In the first paper, binding of the dye rhodamine 6G to the dissociable cyclic trimer glucagon was studied [53]. In this case, ligand fluorescence and absorbance reported on the ligand binding process over different ligand concentration ranges (Fig. 2 and 3 in [53]). The fundamental assumption made in modeling the spectroscopic signals is that absorbance and fluorescence of a bound ligand is independent of both protein oligomerization and the occupancy of other sites on the oligomer (supporting information, Section S4). The signal models, in concert with the dissociable C_3_-NN model, were fitted to the combined experimental data.

The global fit of the model to the data was good (Fig. 5A). As expected, in the absence of any direct data on glucagon oligomerization only one of the equilibrium thermodynamic model parameters (K_olig_) was fully determinable (Fig. 5B). However, from the lower bound on the linkage parameter α_oli_g it is apparent that oligomerization and ligand binding are positively coupled (α_olig_ > 1), and hence the trimer binds ligand more strongly than the monomer. Similarly, from the upper bound on the linkage parameter α_NN_ it is apparent that ligand binding to the oligomer is negatively cooperative (α_NN_ < 1). Interestingly, while α_olig_ is not precisely determined by the data, the product 3*α*_olig_*K*_ligand_is (Fig. S5). This is the basal ligand binding affinity of the trimer (refer Fig. 4).

**Figure 5.**
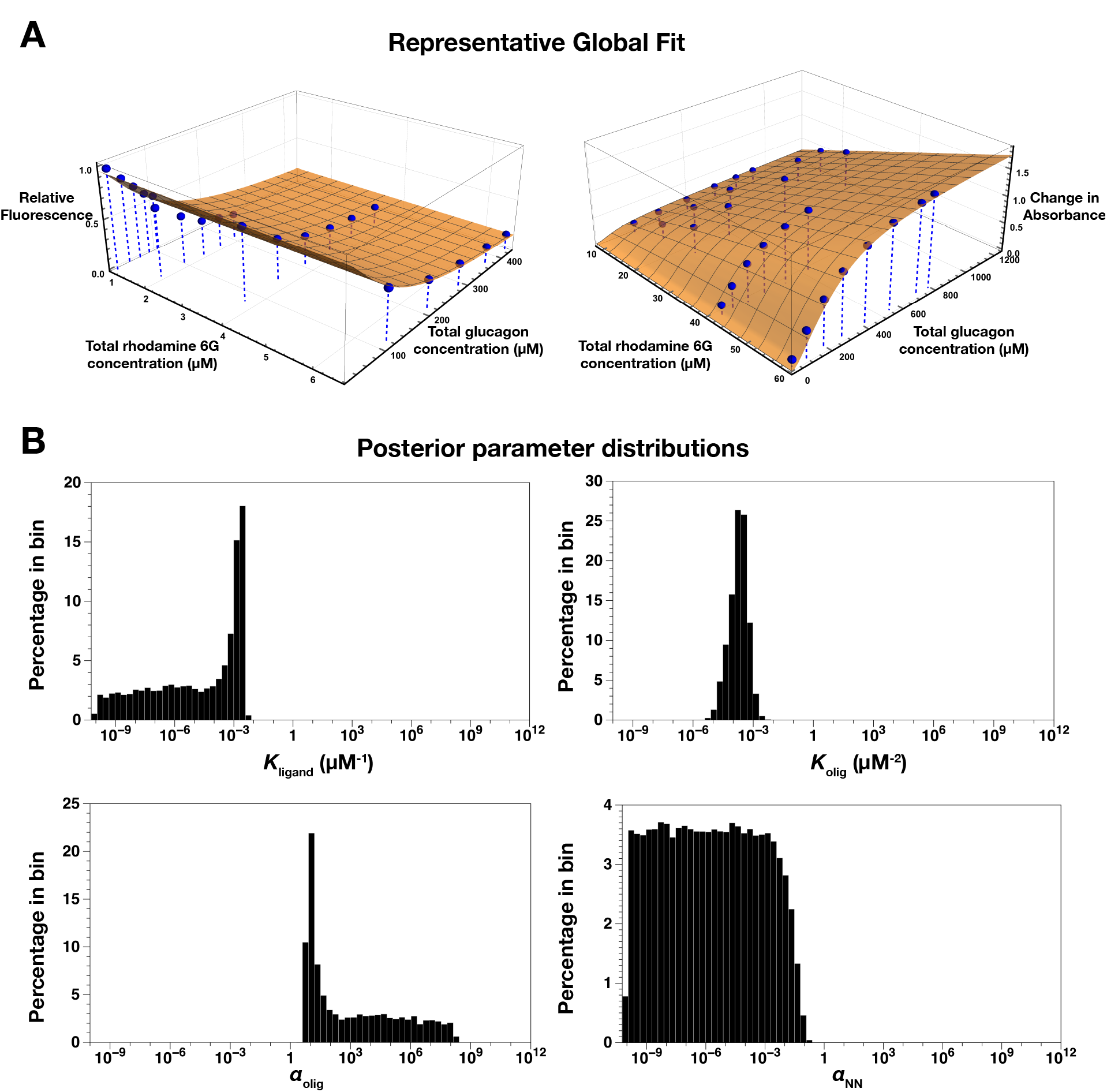
Global fit of the dissociable C_3_-NN model to glucagon data. (A) The blue spheres show the experimental data. Binding of rhodamine 6G was monitored using both fluorescence spectroscopy (data in top left panel) and absorbance spectroscopy (data in top right panel). The orange surfaces show a representative fit of the C_3_-NN model to the data. MCMC analysis was used to fit the model. (B) Posterior distributions of the four parameters of the equilibrium thermodynamic model. Although a wide range of values are consistent with the data, there are sharp cut-offs that effectively indicate upper or lower bounds for the parameter estimates. See also the posterior distribution of the basal ligand association constant of the trimer (Fig S5).

In the original study the authors fitted an equilibrium thermodynamic model that assumed only trimer could bind rhodamine 6G, with a single site per trimer. Fitting of a more general model shows that such restrictive assumptions do not need to be made prior to model fitting, as these linkages are suggested by the data themselves. The revised analysis also better reflects the real uncertainty that exists about the magnitude of the linkage effects.

In the second paper, glucose binding and oligomerization of the dissociable cyclic dimer hexokinase were studied [54]. In this case, intrinsic tryptophan fluorescence and sedimentation equilibria reported on the ligand binding process and the oligomerization process, respectively (Table 4 and Fig. 6 in [54]). The fundamental assumption made in modeling the ligand binding data was that the perturbation of tryptophan fluorescence upon ligand binding to a subunit was independent of both protein oligomerization and the occupancy of the other subunit in the dimer (see supporting information, Section S4). In the case of the oligomerization data, the original analytical centrifugation data were unavailable, so we fit in its place a directly derived quantity - the apparent association constant for the monomer-dimer equilibrium (see supporting information, Section S4). The signal models, in concert with the dissociable C_2_-NN model, were fitted to the combined experimental data.

**Figure 6.**
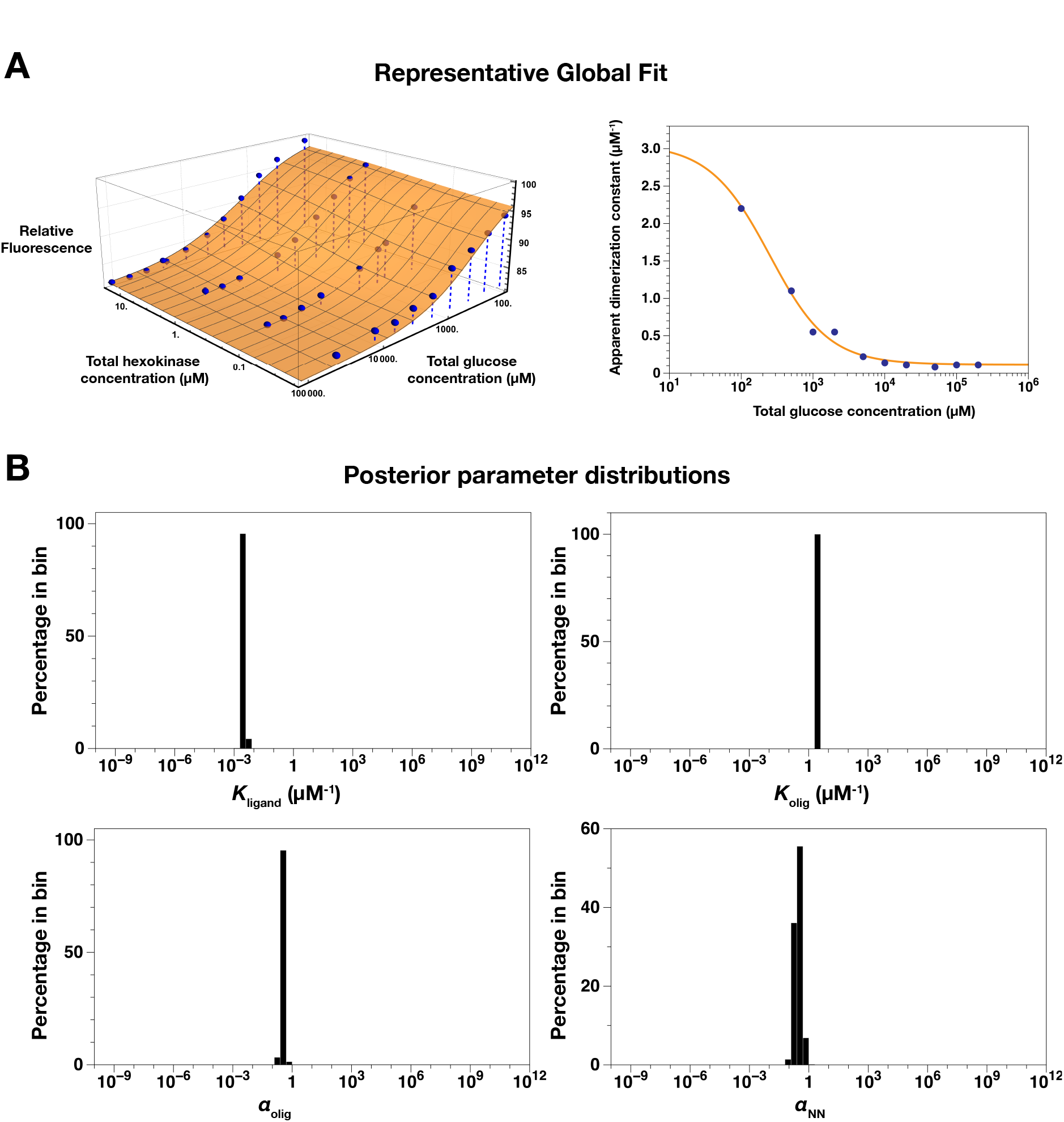
Global fit of the dissociable C_2_-NN model to hexokinase data. (A) The blue spheres/ circles show the experimental data. Binding of glucose was monitored using intrinsic protein fluorescence (data in top left panel). Protein oligomerization was monitored using equilibrium sedimentation analytical ultracentrifugation (derived data in top right panel) The orange surface/line show a representative fit of the C_2_-NN model to the data. MCMC analysis was used to fit the model. (B) Posterior distributions of the four parameters of the equilibrium thermodynamic model. Unlike the fit of the dissociable C_3_-NN model to the glucagon data (Fig. 5), effective point estimates are obtained for each model parameter.

The overall fit of model and data was again good (Fig. 6A). In this case reliable point estimates could be obtained for all parameters of the equilibrium thermodynamic model using the combined ligand binding and oligomerization data (Fig. 6B). Ligand binding appears to exhibit weak negative cooperativity (α_NN_ < 1). Similarly ligand binding and oligomerization are weakly negatively coupled (i.e. binding of glucose promotes dissociation of the hexokinase dimer), a conclusion that was drawn in the original study.

Collectively these results demonstrate the potential of the dissociable C_n_-NN models to determine site-specific thermodynamic information, and quantitate linkage effects, using readily obtained experimental data.

### Theoretical application of the dissociable C_n_-NN models

The dissociable C_n_-NN model parameters explicitly quantitate the thermodynamic couplings that can exist when ligands bind to dissociable ring-like oligomers. When the parameters of the dissociable C_n_-NN models take on extremal values, some interesting and biologically relevant special cases emerge. This idea can be explored using the dissociable C_3_-NN model as an example. The fractional population (f) of protein subunits in each state is given by the individual terms in the binding polynomial, normalized by the total protein subunit concentration:

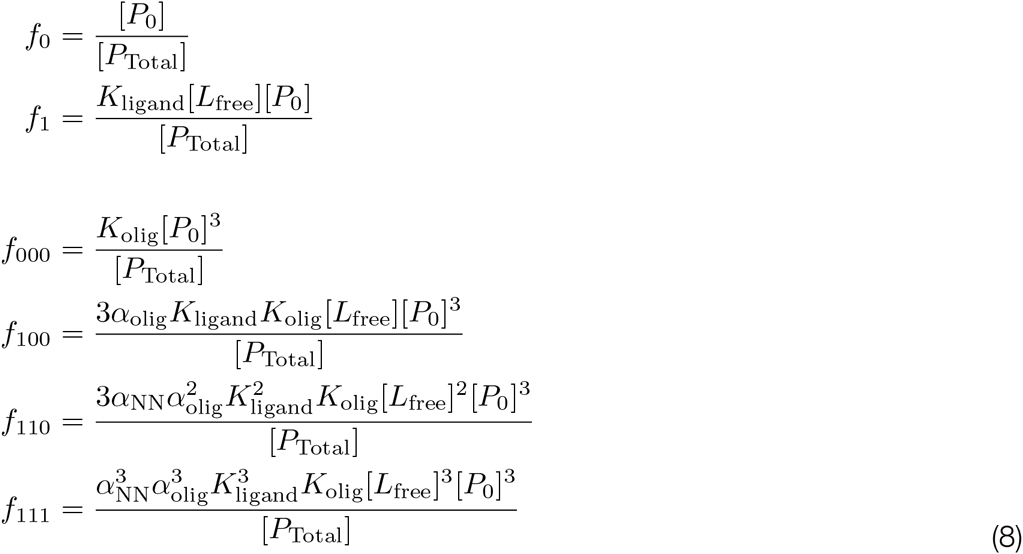

where [*P*_Total_] is given by Equation 6. The limits of these equations can be evaluated as one or more of the association constants or linkage constants goes to zero or infinity, representing extremely unfavorable or extremely favorable binding and linkage, respectively. With limiting values of the model parameters, some of the thermodynamic states become negligibly populated, simplifying the model. We highlight four of the more interesting and commonly encountered special cases that emerge from this analysis. The underpinning mathematics is detailed in the supporting information, Section S6.

Firstly, ligand binding and oligomerization become mutually exclusive (Fig. 7A) when the oligomerization linkage constant becomes very unfavorable (α_olig_ → 0) (Eq. S6.1). This might occur if the ligand binding site and subunit-subunit interface overlap, and the ligand sterically occludes protein association. Secondly, ligand binding becomes obligate for oligomerization (Fig. 7B), when the oligomerization association constant becomes very unfavorable (K_olig_ →0) and the oligomerization linkage constant becomes very favorable (α_olig_ → ∞) (Eq. S6.3b). This might occur if the ligand helps form a major part of the subunit-subunit interface. Thirdly, oligomerization becomes obligate for ligand binding (Fig. 7C), if the ligand association constant becomes very unfavorable (K_ligand_→0) and the oligomerization linkage constant becomes very favorable (α _olig_→∞) (Eq. S6.4b). This might occur if the ligand binding site is created at the subunit-subunit interface, or if the monomer is unstructured and incompetent for ligand binding. Finally, the non-dissociable C*3*-NN model is recovered if the oligomerization association constant is very favorable (K_olig_ → ∞) (Eq. S6.2).

**Figure 7.**
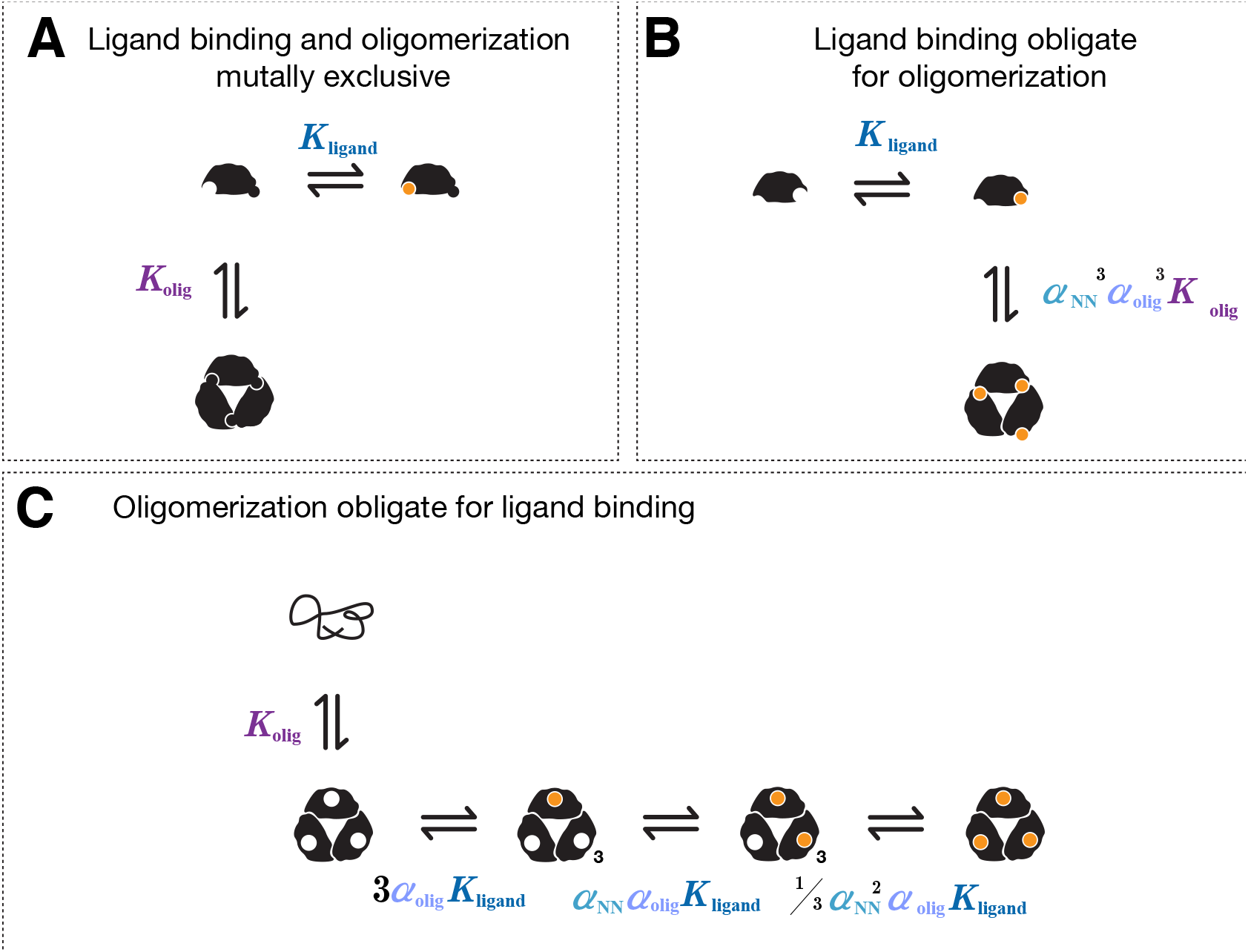
Biologically relevant special cases of the dissociable C_3_-NN model, generated at extremal values of its parameters. (A) Ligand binding and oligomerization become mutually exclusive when the oligomerization linkage constant becomes very unfavorable (*α*_olig_ → 0). This might occur if the ligand binding site and subunit-subunit interface overlap, as depicted. (B) Ligand binding becomes obligate for oligomerization when the oligomerization association constant becomes very unfavorable (*K*_olig_ →0) and the oligomerization linkage constant becomes very favorable (*α*_olig_ → ∞). This might occur if the ligand helps form a major part of the subunit-subunit interface, as depicted. (C) Oligomerization becomes obligate for ligand binding if the ligand association constant becomes very unfavorable (*K*_ligand_ →0) and the oligomerization linkage constant becomes very favorable (*α*_olig_→∞). This might occur if the monomer is unstructured and incompetent for ligand binding, as depicted.

## Discussion

We have developed and analyzed equilibrium thermodynamic models that describe ligand binding to dissociable protein oligomers with cyclic symmetry. These “dissociable C_n_-NN models” (Figs.4 & S3) are site-specific in nature and can account for both linkage between ligand binding events, and linkage between ligand binding and protein oligomerization. We have shown that these models have practical utility, and can be fit to experimental data (Figs. 5 & 6) allowing the linkage effects to be quantified. Furthermore, because of their non-redundant and physically meaningful parameterization, the models are amenable to theoretical analysis. Various biologically relevant special cases arise, when the parameters of the dissociable C_n_-NN models take on extremal values (Fig. 7)

### The Nature and Applicability of the Models

For cyclic protein dimers, which are probably the most ubiquitous system to which the models can be applied, the thermodynamic model is exact, and involves no approximations. For cyclic oligomers of higher order, two key assumptions underpin the models. Firstly, it is assumed that only the monomer and completed cyclic oligomer are significantly populated at equilibrium. The trivial population of intermediates in the assembly and disassembly of cyclic oligomers has been suggested on theoretical grounds [55,56], and a number of experimental studies exist in support [47-49]. Hence this approximation is expected to hold well in most circumstances. Secondly, the models invoke a nearest neighbors approximation, which posits that the energetic consequences of ligand binding are only propagated to the immediate neighbors of each subunit in the ring. This is the central approximation of the Ising model, commonly applied to treat lattice-like systems [44,57], and simply represents a useful and physically reasonable default for the modeling process. If sufficient experimental data were available, and the nearest-neighbors approximation found to be inadequate, additional binary or even ternary interaction parameters could be retained in the model.

By their equilibrium thermodynamic nature, the dissociable C_n_-NN models are blind to the mechanism by which linkage is achieved. Their purpose is simply to detect and quantitate the magnitude of any linkage effects. In this sense they are different to the widely invoked Monod-Wyman-Changeux (MWC) and Koshland-Nemethy-Filmer (KNF) models, which are explicitly mechanistic, and treat cooperative ligand binding effects to protein oligomers as an allosteric phenomenon associated with conformational switching of the protein [Monod: 1965tb; Marzen:2013ir; 58,59]. Because the dissociable C_n_-NN models do not posit a mechanism of linkage, they can be sensibly applied even when the mechanism is unknown. In this way, they are more general than the MWC and KNF models. Subsequently an equilibrium thermodynamic model might be used as the framework on which a fully mechanistic model is developed.

### Model Fitting

The investigation of coupled ligand binding and oligomerization is a relatively complex undertaking, compared to the investigation of simple ligand binding. Not all of the experimental and numerical procedures routinely employed in the simpler case can be used without adaption when studying coupled systems.

Although not our focus in this paper, the model for the experimental signals being measured needs careful consideration. Unlike simple ligand binding systems it is possible that an experimental signal may be perturbed both by ligand binding and by oligomerization. The obvious example is isothermal titration calorimetry, in which the heat released will be dependent on both the change in ligation state and oligomeric state. Spectroscopic signals may be similarly complex. Intrinsic tryptophan fluorescence is frequently used to study both ligand binding and oligomerization in isolation [60]. However when the two processes are coupled, the total fluorescence from the protein may reflect progression of both processes. In such cases the signal model must be correctly constructed, with parameters that account for the perturbation due to each process.

To reliably estimate all parameters of the dissociable C_*n*_-NN model, several types of experimental data are needed, irrespective of their exact nature. Ideally these would report on ligand binding and oligomerization separately, as a function of both total ligand and protein concentration. The experimental conditions (e.g. buffer composition & temperature) need to be the same for all measurements, as the data must be fit globally to the model. When these conditions are satisfied we have shown that all parameters of the dissociable C_*n*_-NN model can be determined uniquely (Fig. 6). However useful information regarding the model parameters can still be obtained from incomplete datasets (Fig. 5) when MCMC methods are employed for parameter estimation.

The use of MCMC methods is suggested because the dissociable C_*n*_-NN models are very non-linear and are fit to multivariate data. Because of these complexities, attempts to minimize the difference between the model and data must account for the many local minima that will be encountered. Related to this, multiple vastly different values of the model parameters may be consistent with the experimental data (Fig. 5). MCMC methods can overcome these problems. In particular the Metropolis-Hastings algorithm can be used to fit an arbitrarily complex model by maximum likelihood and determine the posterior probability distribution of each parameter given the experimental data. This method is preferable to the determination of asymptotic standard errors of parameters via non-linear least squares, which does not generally yield realistic error estimates [50]. Hines has recently provided a useful tutorial in the application of MCMC methods to biophysical problems [24].

Finally, we note that there is generally no closed-form expression for the species concentrations (and hence for the experimental signal) in terms of the dissociable C_*n*_-NN model parameters and the total ligand and protein concentration. This problem can be bypassed by numerical solution of the mass balance equations (Eqns. 4 and 5), with specified values for the thermodynamic model parameters and total concentrations. In practice, the mass balance equations must be numerically solved for each experimental datapoint (i.e. each paired value of total ligand and protein concentration) at each iteration of the algorithm. However the physically relevant solution for the free ligand concentration is bounded by zero and the total ligand concentration, restricting the range for numerical solution. Similar bounds apply to the solution for the unligated protein concentration.

Only in special circumstances can the need for numerical solution of the mass balance equations be bypassed. As noted above, some ligands can be buffered (e.g protons and metal ions), allowing the free ligand concentration to be experimentally controlled. This means that only the protein subunit mass balance equation (Eqn. 4) needs to be solved. For small oligomers (*n* < 5) this allows derivation of closed-form expressions for all species (although the expressions may be very lengthy). The other circumstance in which closed form expressions might occasionally be obtained is when a limiting case of the dissociable C_*n*_-NN model is considered, simplifying the equilibrium thermodynamic model itself.

### Conclusions

The dissociable Cn-NN models developed here should be of utility for quantitative characterization of many proteins. In addition, the systematic derivation of the models and methodology for model fitting and analysis should act as a template for the study of other types of coupled equilibria. Although the computational burden for fitting these models is quite large, with modern computing power and optimized algorithms it is manageable even on personal computers.

## Acknowledgments

We thank Prof. James Sneyd (University of Auckland) for advice on MCMC analysis. MLR was supported by a University of Auckland Doctoral Scholarship.

